# Analysing linear multivariate pattern transformations in neuroimaging data

**DOI:** 10.1101/497180

**Authors:** Alessio Basti, Marieke Mur, Nikolaus Kriegeskorte, Vittorio Pizzella, Laura Marzetti, Olaf Hauk

## Abstract

Most connectivity metrics in neuroimaging research reduce multivariate activity patterns in regions-of-interests (ROIs) to one dimension, which leads to a loss of information. Importantly, it prevents us from investigating the transformations between patterns in different ROIs. Here, we applied linear estimation theory in order to robustly estimate the linear transformations between multivariate fMRI patterns with a cross-validated ridge regression approach. We derived three functional connectivity metrics that describe different features of these voxel-by-voxel mappings: goodness-of-fit, sparsity and pattern deformation. The goodness-of-fit describes the degree to which the patterns in an input region can be described as a linear transformation of patterns in an output region. The sparsity metric, which relies on a Monte Carlo procedure, was introduced in order to test whether the transformation mostly consists of one-to-one mappings between voxels in different regions. Furthermore, we defined a metric for pattern deformation, i.e. the degree to which the transformation rotates or rescales the input patterns. As a proof of concept, we applied these metrics to an event-related fMRI data set consisting of four subjects that has been used in previous studies. We focused on the transformations from early visual cortex (EVC) to inferior temporal cortex (ITC), fusiform face area (FFA) and parahippocampal place area (PPA). Our results suggest that the estimated linear mappings explain a significant amount of response variance in the three output ROIs. The transformation from EVC to ITC shows the highest goodness-of-fit, and those from EVC to FFA and PPA show the expected preference for faces and places as well as animate and inanimate objects, respectively. The pattern transformations are sparse, but sparsity is lower than would have been expected for one-to-one mappings, thus suggesting the presence of one-to-few voxel mappings. The mappings are also characterised by different levels of pattern deformations, thus indicating that the transformations differentially amplify or dampen certain dimensions of the input patterns. While our results are only based on a small number of subjects, they show that our pattern transformation metrics can describe novel aspects of multivariate functional connectivity in neuroimaging data.

**Author summary:** In most functional connectivity studies the multivariate activity patterns associated with regions of interest are reduced to one scalar value per region. This dimensionality reduction approach unavoidably leads to a loss of information and, more importantly, it makes it impossible to investigate the multivariate transformations between patterns of activity in different brain regions. In the current work, we describe methods for estimating the linear transformations between multivariate patterns in pairs of regions, and for characterising their functionally relevant features. In particular, we investigate three main aspects of the mappings: the degree to which they can be considered as linear; the degree to which the transformations can be considered as one-to-one mappings between voxels in the two regions; and the degree to which they geometrically rotate or rescale different input patterns. The experimental findings obtained by applying the novel metrics to an event-related fMRI data set, which reveal the presence of one-to-few voxels mappings between regions involved in different stages of visual processing, clearly show that our methods allow to investigate important and thus far unexplored aspects of multivariate connectivity.

## 1 Introduction

Functional connectivity between brain regions is usually estimated by computing the correlation or coherence between their time series. For this purpose, multivariate (MV) activity patterns within regions of interest (ROIs) are commonly reduced to scalar time series, e.g. by averaging across voxels or by selecting the directions which explain the highest variance (PCA). This process leads to a loss of information and potentially to biased connectivity estimates (Marzetti et al. 2013; Geerligs et al. 2016; Anzellotti et al. 2017, 2018; Basti et al. 2018). Importantly, it also makes it impossible to estimate the transformations between patterns among different ROIs, and to describe functionally relevant features of those mappings. Here, we computed linear MV-pattern transformations between pairs of ROIs in fMRI data, and used them to derive three MV-functional connectivity metrics, i.e. goodness-of-fit, sparsity and pattern deformation.

Recent fMRI studies have explored MV-connectivity between brain regions. For instance, Geerligs et al (2016) applied multivariate distance correlation to resting-state data. This method is sensitive to linear and non-linear dependencies between pattern time courses in two regions of interest. Anzellotti et al. (2016) reduced the dimensionality of their fMRI data per ROI using PCA over time, projecting data for each ROI onto their dominant PCA components. This resulted in a much smaller number of time courses per region than the original number of voxels. Anzellotti et al. (2016) also applied linear (regression) and non-linear (neural network) transformations to the projected lowdimensional data for pairs of brain regions, and found that the non-linear method explained more variance than the linear method. However, dimensionality reduction via PCA leads to a possible loss of information. Indeed, the patterns of the reduced data for different ROIs might not show the same relationships to each other as the original voxel-by-voxel representations. For example, if two regions show a sparse interaction, i.e. each voxel in the first ROI is functionally related only to few voxels in the other ROI, this might not be the case for their corresponding projections on the dominant PCA components. Thus, dimensionality reduction may remove important information about the pattern transformations. Another approach is to ignore the temporal dimension of ROI data and use “representational connectivity”, i.e. compare dissimilarity matrices between two regions (Kriegeskorte et al., 2008a). A dissimilarity matrix describes the intercorrelation of activity patterns for all pairs of stimuli within one region. In this approach, one can test whether the representational structure between two regions is similar or not. However, one cannot test whether the activity patterns of one region are transformations of another, possibly changing the representational structure in a systematic way. Other recent multivariate connectivity approaches also explicitly exploit the presence of functional mappings between regions without characterising the features of these transformations (e.g. Coutanche & Thompson-Schill 2013; Ito et al. 2017).

In the current study, we estimated and analysed the linear transformations between the original voxel-by-voxel patterns. Although it is well-established that transformations of representations between brain areas are non-linear (Naselaris et al. 2011; Khaligh-Razavi & Kriegeskorte 2014; Yamins et al. 2014; Guclu & van Gerven 2015), linear methods can capture a significant amount of the response variance (Anzellotti et al. 2016). Linear transformations are also easy to compute, to visualise, and can be analysed using the vast toolbox of linear algebra. Moreover, our work on linear transformations can serve as a basis for further investigations on MV-connectivity using non-linear transformations. Linear transformations in the case of multivariate connectivity can be described as matrices that are multiplied by patterns of an “input ROI” in order to yield the patterns of an “output ROI”. We can therefore use concepts from linear algebra to describe aspects that are relevant to the functional interpretation of the transformation matrices.

The first concept, similar to the performance metric used in Anzellotti et al. (2016), is that of goodness-of-fit. The degree to which activity patterns in the output region can be explained as a linear transformation of the patterns in the input region is a measure of the strength of functional connectivity between the two regions. The second concept is that of sparsity, i.e. the degree to which a transformation can be described as a one-to-one voxel mapping between input and output regions (a one-to-one voxel correspondence is indeed associated with the highest and non-trivial sparse mapping for each voxel). Topographic maps, in which neighbouring neurons or voxels show similar response characteristics, are well established for sensory brain systems (Patel et al. 2014). It has been suggested that these topographic maps are preserved in connectivity between brain areas, even for higher-level areas (Thivierge & Marcus 2007; Jbabdi et al. 2013). Topography-preserving mappings should result in sparser transformations than those that result in a “smearing” of topographies, or that are random.

Third, we will introduce a measure for pattern deformation. Transformations between brain areas are often assumed to yield different categorisations of stimuli, based on features represented in the output region. The degree to which a transformation is sensitive to different input patterns is reflected in its spectrum of singular values. In the extreme case, where the transformation is only sensitive to one specific type of pattern of the input region but is insensitive to all other orthogonal patterns, it contains only one non-zero singular value. In the other extreme, a transformation which results in a rotation and scaling of all input patterns, would have the maximum number of equal non-zero singular values.

We applied our approach to an existing event-related fMRI data set that has been used in several previous publications to address different conceptual questions (Kriegeskorte et al. 2008a; Kriegeskorte et al. 2008b; Mur et al. 2012; Mur et al. 2013; Jozwik et al. 2016). Four human participants were presented with 96 photographic images of faces (24 images), places (8 images) and objects (64 images). We analysed regions that capture representations at different stages of the ventral visual stream, and that were also the focus of the above-mentioned previous publications, namely early visual cortex (EVC), inferior temporal cortex (ITC), fusiform face area (FFA) and parahippocampal place area (PPA). Specifically, we focused on the transformation from EVC, a region involved at early stages of visual processing, to each of the three other ROIs, which are higher-level regions showing a functional selectivity for the recognition of intact objects (ITC), faces (FFA) and places (PPA) (Malach et al. 1995; Kanwisher et al. 1997; Epstein & Kanwisher 1998).

The aim of our study is to find linear transformations between patterns of beta-values in pairs of ROIs, estimated for different types of stimuli from a general linear model. We here ignored the temporal dimension of the data for two reasons: 1) in fMRI, temporal relationships cannot easily be related to true connectivity unless an explicit biophysical model is assumed; 2) even if such an assumption is made, it would be difficult to estimate a meaningful temporal relationship at the single-trial level as required for this event-related analysis. We therefore focused on spatial pattern information, which in the pre-processing step is estimated from a general linear model. Using this approach, we addressed the following questions (see Fig. 1):

1. To what degree can the functional mappings from EVC to ITC (EVC->ITC), EVC->FFA and EVC->PPA be described as linear matrix transformations? For this purpose, we computed the cross-validated goodness-of-fit of these transformations.
2. To what degree do these transformations represent “one-to-one” mappings between voxels, indicating that they characterise topographical projections? For this purpose, we estimated the sparsity of the transformations.
3. To what degree does a transformation amplify or suppress some MV-patterns more than others? For this purpose, we investigated the degree of pattern deformation by analysing the singular value spectra of the transformations.

**Fig. 1.**
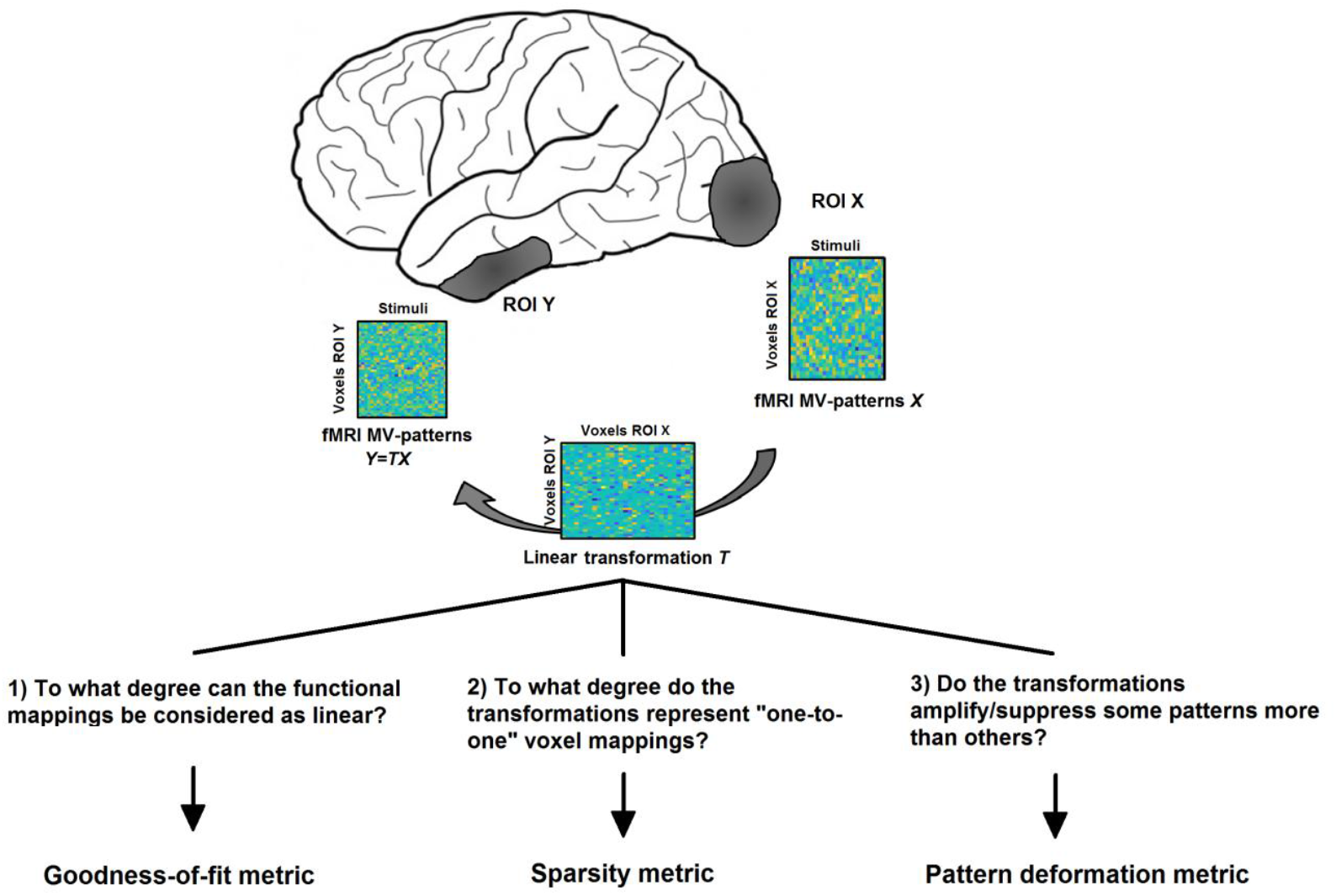
The purpose of the current study is 1) to describe a computational approach for estimating linear transformations ***T*** between MV-pattern matrices *X* and *Y* (each matrix column contains the beta-values associated with a different stimulus type) of two ROIs, and 2) to derive three novel connectivity metrics describing relevant features of those functional mappings.

## 2 Methods

### 2.1 Estimating linear transformations using the ridge regression method

Let us suppose we consider two ROIs X and Y composed of *N_X_* and *N_Y_* voxels, respectively. For each of those two ROIs, we have *N_s_* MV-patterns of beta values obtained from the general linear models with respect to the *N_s_* stimulus types. Let us call the corresponding matrices containing all the MV-patterns ***X*** ∈ *R*^*N_X_*×*N_s_*^ and ***Y*** ∈ *R*^*N_Y_*×*N_s_*^. We also assume that ***X*** and ***Y*** are z-normalised across voxels for each stimulus. We are interested in estimating the transformation ***T*** from ***X*** to ***Y*** and in analysing the features of this transformation. Let us assume that the mapping from ***X*** to the pattern ***Y*** is linear, i.e.

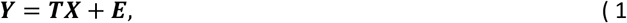

where ***T*** ∈ *R*^*N_Y_*×*N_X_*^ is the transformation matrix and ***E*** ∈ *R*^*N_Y_*×*N_s_*^ is a residual/noise term. The linearity assumption allows us to estimate ***T*** and to investigate its features, e.g. sparsity and singular values. In order to obtain an estimate of the transformation ***T*** we use a ridge regression method (Hoerl & Kennard 1970). Specifically, this method aims to find a suitable solution for ***T*** by minimising the norm of the residuals as well as the norm of the transformation itself. According to this method, the transformation is defined as the matrix

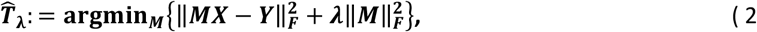

where the parameter *λ* is a positive number which controls the weight of the regularisation term, ***M*** denotes a matrix of the same size of ***T***, and ‖·‖_*F*_ is the matrix Frobenius norm. A unique solution for 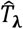 can be obtained using the Moore-Penrose pseudoinverse as

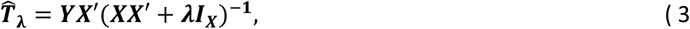

where ***I***_*X*_ ∈ *R*^*N_X_*×*N_X_*^ is the identity matrix and ‘denotes matrix transpose.

#### 2.1.1 Regularisation parameter estimation via cross-validation

Several approaches can be used in order to select a suitable *λ* for ridge regression in eq. (2). These strategies include different cross-validation methods, L-curve and restricted maximum likelihood. Here, we exploit a leave-one-out cross-validation method, which is often used in fMRI studies as a reliable procedure both at stimulus and subject levels (Misaki et al. 2010; Esterman et al. 2010). In our leave-one-stimulus-out procedure, the regularisation parameter is defined as the one which minimises the sum across stimuli of the ratio between the squared norm of the residual and of the (left out) MV-pattern, i.e. as

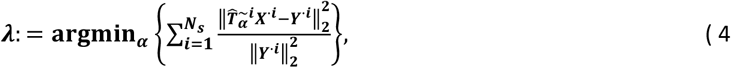

where ***X***^·*i*^ ∈ *R*^*N_X_*×1^ and ***Y***^·*i*^ ∈ *R*^*N_Y_*×1^ are the MV-patterns (beta vectors) associated with the *i*-th stimulus for the two ROIs and 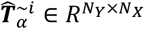 is the transformation matrix obtained by using the MV-patterns of the *N_s_* – 1 stimuli (all the stimuli except for the *i*-th), and with the regularisation parameter α (this approach is nested within the across-sessions cross-validation described in section 2.5.3).

The calculation of the optimal *λ* value would require, for each tested regularisation parameter, the computation of *N_s_* different transformations. However, the computation time can be reduced by using two different observations. First, for demeaned and standardised data, it holds that 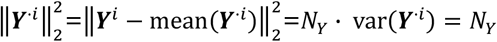. We can thus rewrite the previous formulation for *λ* without considering the denominator within the sum, i.e. the value of *λ* can be now obtained by minimising the sum of squared residuals. Secondly, as is shown in Golub et al. (1979), the *λ* value obtained in such a way is equivalent to that obtained by minimising the functional 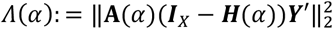, where **A**(*α*) is the diagonal matrix whose non-zero entries are equal to 1/(1 – *h_ii_*(*α*)), being the *h_ii_*(*α*) the *ii*-th elements of ***H***(*α*):= ***X*′(*XX*″** + *α**I**_X_*)^−1^***X***. Using the two previous observations, we can finally assess the value of *λ* as

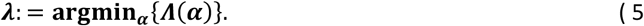

This final formulation allows us to obtain the optimal value in a reduced computation time, thus also facilitating the calculation of the goodness-of-fit metric (see below).

### 2.2 Characterisation of the goodness-of-fit

In order to assess the goodness-of-fit of the MV-pattern transformations between ROIs, we compute the cross-validated percentage of pattern variance in the output region which can be explained using a linear transformation of patterns in the input region, with an optimal regularisation parameter *λ* obtained as above. Specifically, we define the percentage goodness-of-fit (*GOF*) as

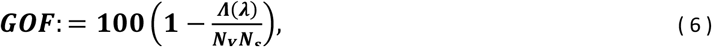

where *Λ*(·) is the functional which describes the sum of squared residuals (see section 2.1.1). This metric can be considered as a method to quantify the (linear) statistical dependencies among the MV-patterns by means of the explained output pattern variance. A GOF value equal to 100 denotes a perfect linear mapping while low values suggest that either the mapping does not exist or it is fully non-linear.

To assess the statistical significance of the observed GOF values, we exploit a permutation test (with 10,000 permutations of the EVC patterns, in order to randomise the stimuli). We compare the distribution consisting of the GOF values for the four subjects with the reference distribution obtained from permutation, relying on the Kolmogorov-Smirnov (K-S) test. We consider the observed GOF as significant when the *p*-value is lower than 0.05.

### 2.3 Characterisation of the transformation sparsity

A sparse matrix is defined as having the majority of its elements equal to 0 (Stoer & Bulirsch, 2002). In the case of MV-pattern transformations between ROIs, a sparse matrix could indicate a one-to-one mapping between voxels in the two ROIs. However, even in the presence of a perfect sparse linear mapping, we cannot expect the elements of the estimated 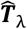 to be exactly zero (Fig. 2, panels A and B), because the ridge regression method always leads to a smooth solution. However, other approaches, such as the least absolute shrinkage and selection operator (LASSO, Tibshirani 1996), may lead to the opposite problem, i.e. obtaining sparse solutions even in the presence of non-sparse linear mappings (see Fig. S1). We therefore need to define a strategy to reliably estimate the degree of sparsity of the transformation matrix.

The idea behind our approach is to take into account both the GOF value, taken as a measure of the level of noise (i.e. the higher the GOF value the lower the noise level), and the rate of decay of the *density* curve. The *density* curve describes the fraction of the entries of the estimated transformation which are larger than a threshold, as a function of this threshold. The steepness of the decay of this curve increases with the increase of the degree of sparsity of the original transformation, and it decreases with the increase of the level of noise in the model.

**Fig. 2.**
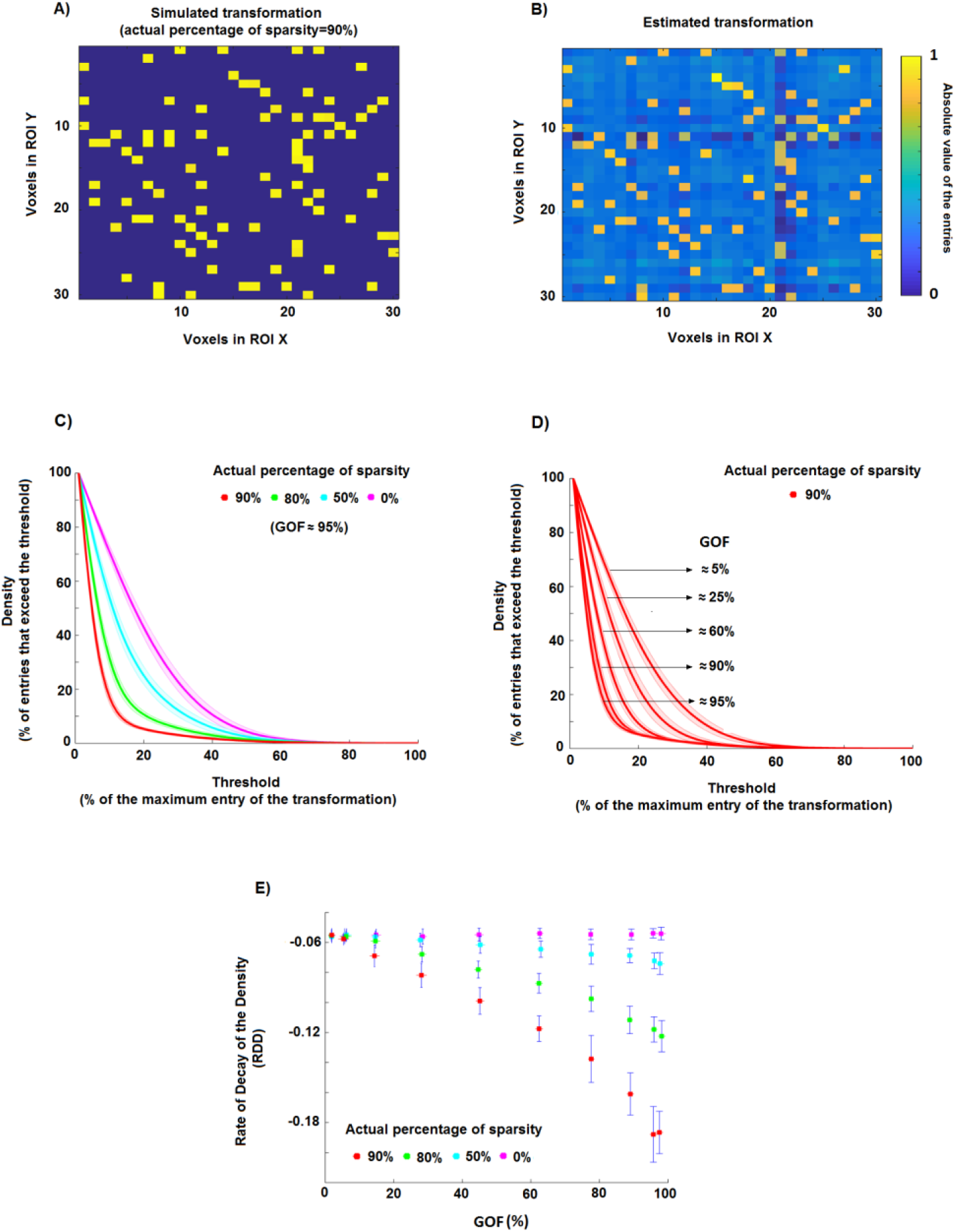
Estimation of the percentage of sparsity using a ridge regression method. **A)** An example of a simple sparse simulated transformation: 90% of the entries are equal to 0, while the other 10% of the entries are equal to 1. **B)** Estimate of the transformation in A obtained by using the ridge regression method. The lighter background indicates that the estimated elements are different from zero even if in the original transformations they are exactly equal to zero. **C)** Density of the thresholded estimated transformations, i.e. the percentage of matrix entries that exceed the threshold, as a function of the threshold. In this toy example, we generated 30 realisations for four simulated percentages of sparsity (0%, 50%, 80% and 90%). **D)** Density of the thresholded estimated transformations associated with a degree of sparsity of 90% and different GOF values (i.e. different levels of noise in the model). **E)** Scatter plots of the rate of decay of the density curves (*RDD*) shown in panel C against goodness-of-fit *GOF* for the 30 simulation realisations of each of the four different cases. It is evident that simulated transformations of e.g. 90% sparsity are associated with a certain range of *RDD* and *GOF* values (red dots) which, at least for sufficiently large values of *GOF*, are different from those related to transformations with 80% of sparsity (green dots).

Let us take the normalised 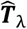 obtained by dividing its elements by its maximum absolute value. We define the density curve *d* as the function of the threshold *P* ∈ [0,1] which describes the fraction of elements of 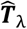 whose absolute value exceeds *P*. Specifically, *d* is defined as

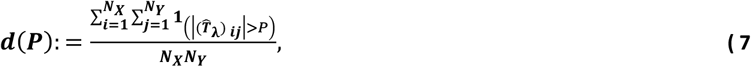

where 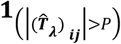 denotes the indicator function of the set 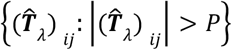, which is equal to 1 if 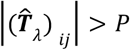 holds, and 0 otherwise. The density curve *d* is a monotonically decreasing function of *P*, with a value of 1 for *P* = 0 and of 0 for *P* = 1.

The analysis of the rate of decay of the density curve of 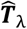 as a function of the threshold *P* provides an estimate of the actual degree of sparsity of ***T***. Higher degrees of sparsity are associated with steeper decay. For instance, panel C of Fig. 2 shows the *d* curve for five different toy cases in a noise free situation. For each case, we simulated 30 multivariate patterns ***X*** (of size 128 voxels x 96 stimuli) and 30 transformations ***T*** (128 voxels x 128 voxels) as following standard normal distributions. Each of the five cases in this toy example has a different true percentage of sparsity, i.e. 0%, 50%, 80%, 90%, obtained by randomly setting to 0 the corresponding percentage of elements of ***T***. We then calculate the MV-pattern matrix as ***Y*** as ***Y*** = ***TX***. The density curves (average and standard deviation across 30 realisations for each case are denoted by solid lines and shaded areas) are clearly distinguishable from each other, thus allowing us to disentangle the five cases.

However, the rate with which a density curve decays from 1 to 0 depends on the noise level in the model. In particular, the steepness of the decay associated with a fixed degree of sparsity decreases with the increase of the level of noise, i.e. with the decrease of the GOF value (panel D, Fig. 2). Thus, by only analysing the density curve it is not possible to distinguish two different percentages of sparsity for which the noise levels are different, i.e. the curve *d* for a percentage of sparsity *S*_1_ with noise level *L*_1_ can be undistinguishable from the curve for a sparsity *S*_2_ with noise level *L*_2_. To overcome this problem, we take into account both the density curve *d* and the goodness-of-fit *GOF* between the MV-patterns. As shown in the panel E of Fig. 2, by using 1) the rate of decay of the density curve (*RDD*), defined as the parameter *b* of an exponential function *aexp*(*bP*) fitted to the *d* function with a non-linear least squares fitting method, and 2) the value of *GOF*, it is possible to disentangle the simulated degree of sparsity even if the noise levels are different. Let us now describe step by step the Monte Carlo based approach that we use in order to estimate the degree of sparsity of the pattern transformations in our data set. We consider the transformations EVC->ITC, EVC->FFA and EVC->PPA for the set of stimuli composed of the 96 images of all stimulus types.

#### 2.3.1 Monte Carlo approach to obtain the percentage of sparsity

Let us suppose we are interested in estimating the degree of sparsity of the pattern transformation between EVC (consisting of 224 voxels in our data) and ITC (316 voxels) by using the patterns obtained from the full set of 96 stimuli. The other cases will be analogously treated. The strategy described below can be considered as a Monte Carlo method. Specifically, we:

1. simulate, for each noise level and each percentage of sparsity, transformations ***T*** (size 256 voxels x 224 voxels), the non-zero entries of which follow a standard normal distribution and the positions of the zero entries were randomly selected;
2. compute estimates 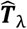 of each true ***T*** by using a ridge regression method on the original EVC patterns ***X*** and the simulated ITC pattern 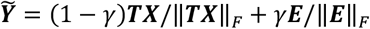 (all the patterns were first demeaned and standardised), where ***E*** and *γ* denote the independent Gaussian noise/residual signal and its relative strength. The estimated transformation is given by

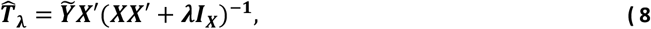

where the *λ* value denotes the same regularisation parameter obtained on real data via the cross-validation procedure described in section 2.1.1. Importantly, the characteristics of the simulated data were chosen to resemble those of the real MV-patterns. The patterns in the real data follow a standard normal distribution: none of the 3072 patterns (4 regions, 4 subjects, 2 sessions and 96 stimuli) deviate significantly from normality as assessed by a one-sample Kolmogorov-Smirnov test with the null hypothesis that the patterns follow a Gaussian distribution with zero mean and a standard deviation equal to one (*p* > 0.05, Bonferroni-corrected for multiple comparisons). In Fig. S2 we provide, as an example, the histogram of the values of the actual (left panel) and simulated (right panel) MV-patterns of the ITC for one subject.
3. calculate, for each 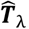, the density curve *d*, its rate of decay *RDD*, and the goodness-of-fit *GOF*;
4. calculate the average *RDD* and *GOF* across the simulation-realisations for each different simulated percentage of sparsity and noise level. In such a way, we obtain, for each simulated percentage of sparsity, a curve describing the mean *RDD* value as a function of the *GOF*.
5. estimate the percentage of sparsity for the real data by looking at the point of coordinates equal to the average (across subjects) *RDD* and *GOF*. For instance, if this point lies between two curves representing the results for 50% and 60% of sparsity, the estimated sparsity of the transformation EVC->ITC would be 50-60%.

We simulate six different percentages of sparsity, i.e. 50%, 60%, 70%, 80%, 90% and 99%. An estimated percentage of sparsity lower than 50% indicates that the transformation is not sparse (the majority of elements is different from 0) while a higher value in the simulated range indicates the opposite. We use 10 different levels of noise strength: the γ value ranged from 0 to 0.9 with a step of 0.1. For each different percentage of sparsity and noise level, the number of simulations is 1000.

### 2.4 Characterisation of the induced pattern deformation

A MV-pattern of each ROI can be considered as a point belonging to a vector space whose dimension is equal to the number of voxels in that region. A MV-pattern transformation thus corresponds to the linear mapping ***T***:*R^N_X_^*→*R^N_Y_^* between the two respective vector spaces. The aim of this section is to: 1) explain why the singular values (SVs) of the transformation 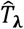 are important features of this mapping, and 2) describe the computational strategy used to understand how much the transformation deformed the original patterns, e.g. via asymmetric amplifications or compressions along specific directions.

Let us assume for simplicity’s sake that *N_X_* is equal to *N_Y_*, i.e. that the number of voxels in the ROI X is equal to the number of voxels in the ROI Y. By means of the polar decomposition theorem (Nigham 1986) we can consider the transformation ***T*** as the composition of an orthogonal matrix ***R*** multiplied by a symmetric positive-semidefinite matrix ***P***_1_ or as the composition of a different symmetric positive-semidefinite matrix ***P***_2_ followed by the same matrix ***R***, i.e.

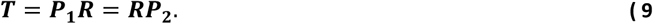

This factorisation has a useful heuristic interpretation (panel A of Fig. 4). It states that can be written in terms of simple rotation/reflection (i.e. the matrix ***R***) and scaling pattern transformations (i.e. the matrices ***P***_1_ and ***P***_2_). Furthermore, even if the square matrix ***T*** is not a full rank matrix, ***P***_1_ and ***P***_2_ are unique and respectively equal to 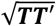 and 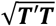, where ‘denotes the transpose. It is also evident that the eigenvalues of ***P***_1_ and ***P***_2_, which indicate the scaling deformation factors induced by ***T***, are equal between the two and coincide with the SVs of the pattern transformation ***T*** (Nigham 1986). Although, for the sake of clarity, we discussed the geometrical framework explicitly assuming the two ROIs to be composed of the same number of voxels, a similar argument can be made for the general case. In other words, the singular values of the transformation are important features of the mapping that describe the “dampening” or “amplification” of different dimensions even if the sizes of the two ROIs are different.

In order to investigate the pattern deformations, we analysed the SVs of the estimated transformation 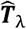 (panel B of Fig. 4 shows estimates of SVs for some simulated toy examples). For this purpose, we defined a metric describing the average pattern deformation induced by the transformation ***T***. We computed the rate of decay of the SVs of the estimated transformation 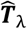 (*RDSV*), defined as the parameter *b* of an exponential function fitted to the curve composed of all the SVs (as in 2.3 for the *d* curve). For instance, a value of 0 for *RDSV* corresponds to constant values for the SVs, i.e. the mapping induces the same deformation between the MV-patterns associated with each stimulus, while a larger *RDSV* value is associated with a larger asymmetric deformation, i.e. the patterns are differently amplified/compressed before or after rotation depending on the stimulus.

Additionally, the rate with which the SVs of 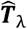 decay depends on the degree to which the MV-patterns in the output region can be described by the linear mapping from the input region (panel C, Fig. 4). Therefore, as for the previously described strategy for characterising the sparsity of the transformations, we also take into account the goodness-of-fit *GOF* between the patterns as a measure of the level of noise in the model (panel D of Fig. 4). In this way, we can understand if two transformations induce a different deformation on the MV-patterns in the presence of different levels of noise. The SVs of the estimated transformations can also be exploited for investigating the pattern deformation in the general case in which there is a different number of voxels in the two ROIs. Let us now describe step by step the Monte Carlo based approach that we use in order to estimate the average pattern deformation induced by the transformations between EVC and the other three ROIs (i.e., EVC->ITC, EVC->FFA and EVC->PPA).

#### 2.4.1 Monte Carlo approach to obtain the rate of decay of singular values curve

Let us suppose we are interested in investigating the pattern deformation for the same transformation for which we assessed sparsity in 2.3.1, i.e. between EVC and ITC, using the MV-patterns obtained from the full set of 96 stimuli. The other cases will be analogously treated. The Monte Carlo approach that we use consists of the following steps:

1. we simulate, for each noise level and each rate of exponential decay of the SVs, transformations ***T***. For each realisation of ***T***:

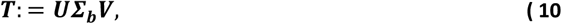

where ***U*** and ***V*** are two orthogonal matrices obtained by applying a singular value decomposition on a matrix whose entries follow standard Normal distributions, and Σ_*b*_ is a diagonal matrix whose non-zero entries follow an exponential decay with parameter *b*;
2. as in the section 2.3.1, we compute the estimates 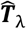 of the true ***T*** by using a ridge regression method on the original EVC patterns ***X*** and the simulated ITC pattern obtained as 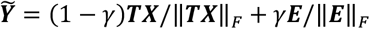 (the patterns were first demeaned and standardised for each stimulus);
3. we calculate the *RDSV*, i.e. the rate of decay of the SVs for the estimated transformation (we only use the *P* largest values with *P* = *m* in{ rank (*X*), rank (*Y*)) }, and the goodness-of-fit *GOF*;
4. we calculate the average *RDSV* and *GOF* across the simulation-realisations for each different simulated decay of the SV-curve and noise level. In such a way, we obtain, for each simulated decay, a curve describing the mean *RDSV* value as a function of the *GOF*;
5. we estimate the pattern deformation for the real data by looking in the *RDSV* – *GOF* plane at the point of coordinates equal to the average (across subjects) *RDSV* and *GOF*. For example, if this point lies between two curves representing the results for the rates of decay of *b* = −0.0 1 and *b* = 0, the estimated decay of the SVs curve would be (−0.0 1, 0).

**Fig. 3.**
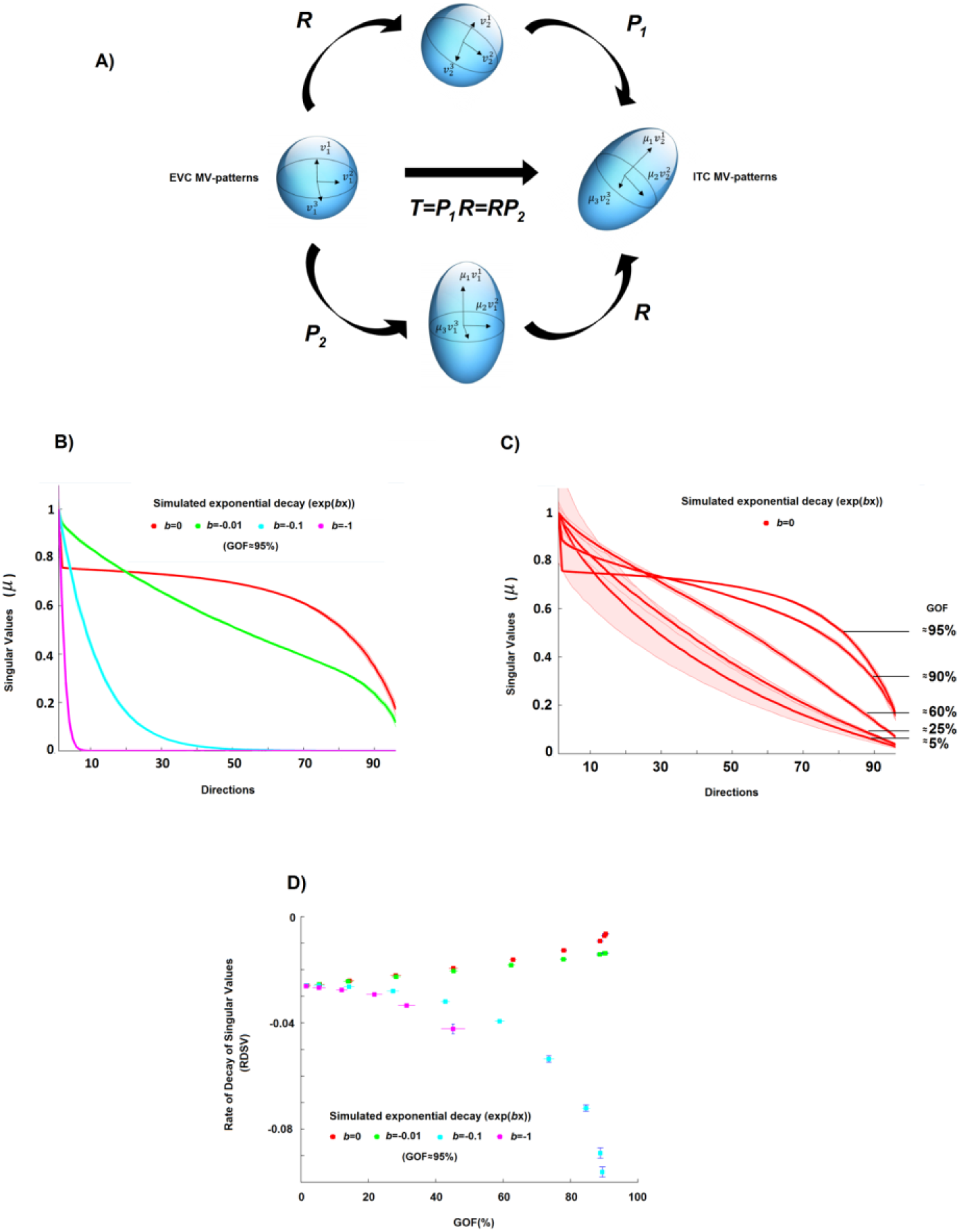
Estimation of the pattern deformation. **A)** A geometric interpretation of a linear pattern transformation between patterns of equal dimension. Two MV-patterns of two ROIs, let us say EVC and ITC, can be seen as two points of two vector spaces, and the matrix transformation ***T*** between them can be seen as a linear mapping between these two vector spaces. In this panel, a sphere (representing for simplicity the MV-patterns of EVC for a set of stimuli) is transformed by *T* into the ellipsoid (representing the ITC MV-patterns for the same set of stimuli). The singular values (SVs) of *T* are important features of this mapping. For example, if the number of voxels is the same in both ROIs, the SVs (*μ* values in the figure) describe how much the EVC pattern is deformed by the transformation. For instance, constant values across all SVs can indicate an orthogonal transformation, that is, a linear mapping in which the ITC pattern can be completely described as a rotation (or reflection) of the original EVC pattern. **B)** The curves of the SVs (a monotonically non-increasing function with *b* as the rate of decay) of the estimated transformations for four different simulated rates of decay (*b* = 0, −0.01, −0.1, −1). **C)** The curves of the SVs of the estimated transformations associated with orthogonal transformations (i.e., *b* = 0) and different GOF values (i.e. different levels of noise in the model). **D)** Scatter plot between goodness-of-fit *GOF* and the rate of decay of singular values *RDSV*, i.e. the estimated decay obtained by fitting an exponential curve to the SVs of the estimated transformation for the four different cases. By using both the *RDSV* and *GOF*, it is possible to characterise the different induced pattern deformations, even if the level of noise is not equal to 0%.

We use four different simulated rates of exponential decay of the SVs, which indicate four different orders of magnitude of the exponential decay: 0, −0.0 1, −0.1, −1. We also use 10 different levels of noise strength γ, which ranged from 0 to 0.9 with a step of 0.1. For each different rate of exponential decay and level of noise, the number of simulations is 1000.

### 2.5 Real fMRI data

The fMRI data set has been used in previous publications (Kriegeskorte et al., 2008a; Kriegeskorte, et al., 2008b; Mur et al., 2012; Mur et al. 2013; Jozwik et al. 2016). Four healthy human volunteers participated in the fMRI experiment (mean age 35 years; two females).

The stimuli were 96 colour photographs (175 x 175 pixels) of isolated real-world objects displayed on a gray background. The objects included natural and artificial inanimate objects as well as faces (24 photographs), bodies of humans and nonhuman animals, and places (8 photographs). Stimuli were displayed at 2.9° of visual angle and presented using a rapid event-related design (stimulus duration: 300 ms, interstimulus interval: 3700 ms) while subjects performed a fixationcross-colour detection task. Each of the 96 object images was presented once per run in random order. Subjects participated in two sessions of six 9-min runs each. The sessions were acquired on separate days.

Subjects participated in an independent block design experiment that was designed to localise regions of interest (ROIs). The block-localiser experiment used the same fMRI sequence as the 96 images experiment and a separate set of stimuli. Stimuli were grayscale photos of faces, objects, and places, displayed at a width of 5.7° of visual angle, centered with respect to a fixation cross. The photos were presented in 30 s category blocks (stimulus duration: 700 ms, interstimulus interval: 300 ms), intermixed with 20 s fixation blocks, for a total run time of 8 min. Subjects performed a one-back repetition detection task on the images.

#### 2.5.1 Acquisition and Analysis

##### Acquisition

Blood oxygen level-dependent (BOLD) fMRI measurements were performed at high spatial resolution (voxel volume: 1.95 × 1.95 × 2mm^3^), using a 3 T General Electric HDx MRI scanner, and a custom-made 16-channel head coil (Nova Medical). Single-shot gradient-recalled echo-planar imaging with sensitivity encoding (matrix size: 128 × 96, TR: 2 s, TE: 30 ms, 272 volumes per run) was used to acquire 25 axial slices that covered inferior temporal cortex (ITC) and early visual cortex (EVC) bilaterally.

##### Pre-processing

fMRI data preprocessing was performed using BrainVoyager QX 1.8 (Brain Innovation). All functional runs were subjected to slice-scan-time correction and 3D motion correction. In addition, the localiser runs were high-pass filtered in the temporal domain with a filter of two cycles per run (corresponding to a cutoff frequency of 0.004 Hz). For the definition of FFA and PPA (see 2.5.2 below), data were spatially smoothed by convolution of a Gaussian kernel of 4 mm full-width at half-maximum. For definition of EVC and ITC, unsmoothed data were used. Data were converted to percentage signal change. Analyses were performed in native subject space (i.e., no Talairach transformation).

##### Estimation of single-image patterns

Single-image BOLD fMRI activation was estimated by univariate linear modeling. We concatenated the runs within a session along the temporal dimension. For each ROI, data were extracted. We then performed univariate linear modelling for each voxel in each ROI to obtain response-amplitude estimates for each of the 96 stimuli. The model included a hemodynamic-response predictor for each of the 96 stimuli. The predictor time courses were computed using a linear model of the hemodynamic response (Boynton et al., 1996) and assuming an instant-onset rectangular neuronal response during each condition of visual stimulation. For each run, the design matrix included the stimulus-response predictors along with six head-motion parameter time courses, a linear-trend predictor, a six-predictor Fourier basis for nonlinear trends (sines and cosines of up to three cycles per run), and a confound-mean predictor.

#### 2.5.2 ROI definition

ROIs were defined based on visual responsiveness (for EVC and ITC) and category-selective contrasts (for fusiform face area, FFA, and parahippocampal place area, PPA) of voxels in the independent block-localiser task and restricted to a cortex mask manually drawn on each subject’s fMRI slices (Mur et al., 2012).

The FFA was defined in each hemisphere as a cluster of contiguous face-selective voxels in ITC cortex (number of voxels per hemisphere: 128). Face-selectivity was assessed by the contrast faces minus places and objects.

Clusters were obtained separately in the left and right hemisphere by selecting the peak face-selective voxel in the fusiform gyrus, and then growing the region from this seed by an iterative process. During this iterative process, the region is grown one voxel at a time, until an a priori specified number of voxels is selected. The region is grown by repeatedly adding the most face-selective voxel from the voxels that are directly adjacent to the current ROI in 3D space, i.e., from those voxels that are on the “fringe” of the current ROI (the current ROI is equivalent to the seed voxel during the first iteration).

The PPA was defined in an identical way but then using the contrast places minus faces and objects, growing the region from the peak place-selective voxel in the parahippocampal cortex in each hemisphere (number of voxels per hemisphere: 128).

The ITC ROI was defined by selecting the most visually responsive voxels within the ITC portion of the cortex mask (number of voxels for the bilateral ITC region: 316). Visual responsiveness was assessed by the contrast visual stimulation (face, object, place) minus baseline.

In order to define EVC, we selected the most visually responsive voxels, as for ITC, but within a manually defined anatomical region around the calcarine sulci within the bilateral cortex mask (number of voxels: 224). For EVC and ITC, voxels were not constrained to be spatially contiguous.

#### 2.5.3 Metrics calculation on real data

In order to estimate and analyse the transformation between the input MV-patterns of EVC and the output MV-patterns of ITC, FFA and PPA, we relied on an across-sessions approach. We first estimated the transformation 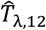, as well as the values of *RDD*_12_ and *RDSV*_12_ the cross-validated *GOF*_12_, from the input patterns of session 1 and the output patterns of session 2. Second, we estimated 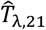 and the values of the three metrics, by using the input/output patterns of the session 2/1. Then, we averaged the obtained values, i.e. we defined *GOF*:=(*GOF*_2_ + *GOF*_21_)/2, *RDD*: = (*RDD*_12_ + *RDD*_21_)/2 and *RDSV*: = (*RDD*_12_ + *RDSV*_12_)/2. The application of an across-sessions approach improves the interpretability of the measures by reducing possible confounds induced by stimulus-unrelated intrinsic fluctuations shared between brain regions (Henriksson et al. 2015; Walther et al. 2016).

## 3 Results

### 3.1 Goodness-of-fit, explained variance

The results obtained by analysing the goodness-of-fit (*GOF*, panel A, Fig. 4) associated with the transformations (Fig. S3 show the estimated mappings for each subject and pair of ROIs as heat maps) clearly show the presence of a linear statistical dependency between EVC and ITC, FFA and PPA. The cross-validated average *GOF* value across the four subjects for each pair of ROIs is statistically different from the *GOF* values obtained by permutation test (*p* < 0.05, K-S test). The GOF value achieved by EVC in estimating the ITC patterns shows the largest value. The EVC->ITC value is significantly larger (*p*= 0.001, paired t-test, Cohen’s d = 5.97) than the value associated with EVC->PPA. No statistically significant difference can be observed between EVC->ITC and EVC->FFA (*p*=0.066, paired t-test, Cohen’s d = 1.41), as well as between EVC->FFA and EVC->PPA (p=0.069, paired-test, Cohen’s d = 1.39). In order to show the reliability of the GOF metric at a within subject level, we compared the session1-session2 transformations to the session2-session1 mappings. We found that the goodness-of-fit (GOF) shows similar values across sessions: 26.9 6.4 and 27.8 7.6 for EVC-ITC; 14.2 ± 3.7 and 16.8 ± 5.9 for EVC-FFA; 6.6 ± 4.3 and 8.8 ± 4.2 for EVC-PPA.

**Fig. 4.**
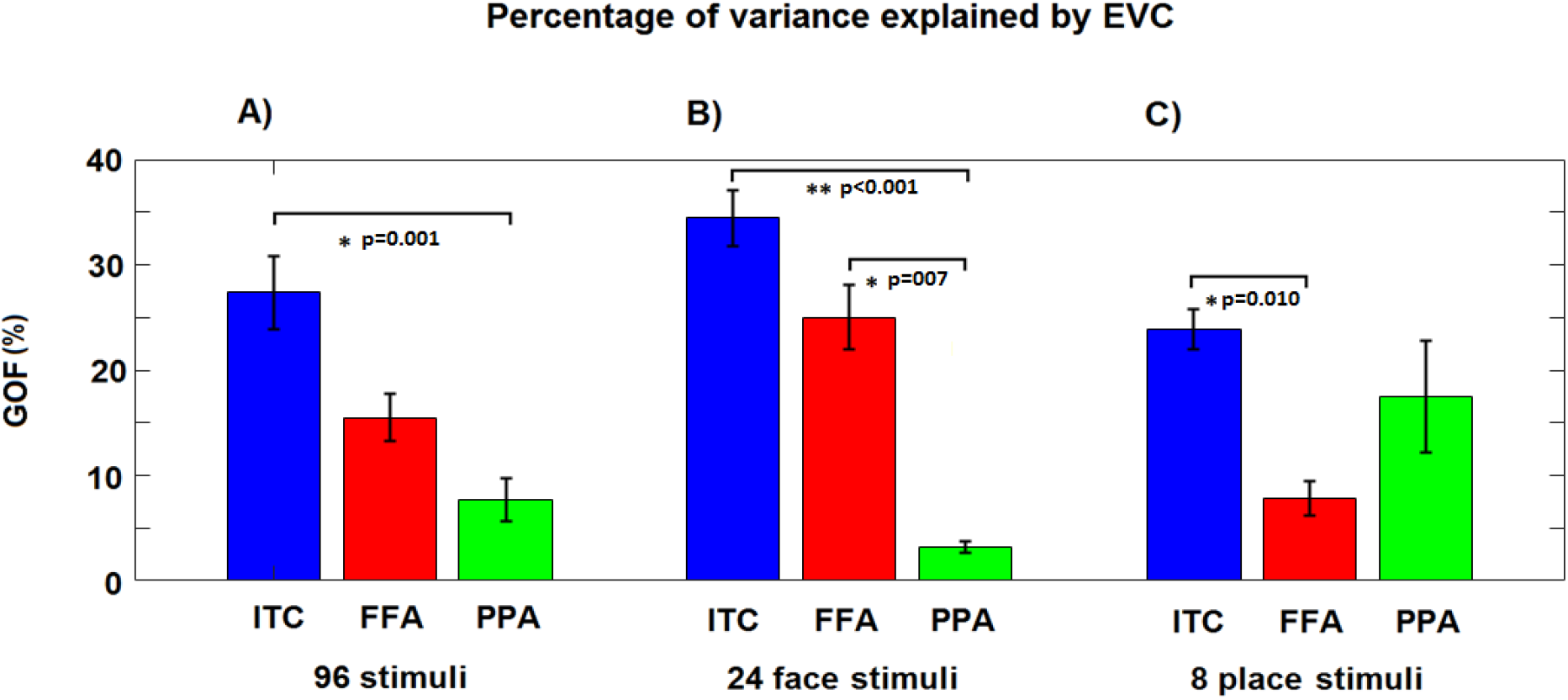
The goodness-of-fit (*GOF*) values for the three sets of stimuli. **A), B)** and **C)** The average (and standard error) percentages of *GOF* by using the linear pattern transformations from EVC to the three output ROIs for all 96 stimuli, the 24 face stimuli and the 8 place stimuli, respectively. The *p*-values for the paired *t*-tests are reported even if based on only four subjects.

We then separately ran two analyses that, instead of considering all the 96 stimuli, only considered the 24 face and 8 place stimuli (which are included in the set composed of the 96 original stimuli). Middle and right panels of Fig. 4 show the results. For the subset of stimuli composed of faces (panel B, Fig. 4), the GOF value for EVC->ITC is significantly larger (*p* < 0.001, paired t-test, Cohen’s d = 7.41) than the GOF value associated with EVC->PPA. The percentage of variance of FFA explained by EVC (i.e. EVC->FFA) is significantly (*p*=0.007, paired t-test, Cohen’s d = 3.31) larger than that for EVC->PPA, while no statistically significant difference can be observed between EVC->ITC and EVC->FFA (p=0.113, paired t-test, Cohen’s d= 1.06). For the subset of 8 places stimuli (panel C, Fig. 4), a significant difference (*p*=0.010, paired t-test, Cohen’s d = 2.95) can only be observed EVC->ITC and EVC->FFA. No statistically significant differences can be observed between EVC->FFA and EVC->PPA (*p*=0.157, paired t-test, Cohen’s d=0.9 4) and between EVC->ITC and EVC->PPA (*p*=0.371, paired t-test, Cohen’s d = 0.52).

Fig. S4 shows the *GOF* as a function of the stimulus. The first 48 stimuli are images of animate objects (including animal and human faces) while the last 48 stimuli are images of inanimate objects (including natural and artificial places). For ITC and FFA, the *GOF* is generally higher for the animate than inanimate objects, while the opposite is observed for PPA. For 39 of the 48 animate objects, the *GOF* for EVC->PPA is lower than the average *GOF* across the 96 stimuli, while for 35 of the 48 inanimate objects, the *GOF* is higher (*p*<0.001, Fisher exact test). Conversely, for 38 of the 48 animate objects, the *GOF* for EVC->FFA is higher than the average *GOF* across the 96 stimuli, while for 36 of the 48 inanimate objects, the *GOF* is lower (*p*<0.001, Fisher exact test). The results for EVC->ITC are similar to those for EVC->FFA: for 35 of the 48 animate objects, the *GOF* for EVC->ITC is higher than average, while for 27 of the 48 inanimate objects, the *GOF* is lower (*p*<0.01, Fisher exact test).

As an alternative to explained variance (i.e. to GOF metric), it is possible to analyse the correlation between the dissimilarity matrix associated with a pattern and the matrix obtained using the output pattern 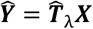. Fig. 5 shows the dissimilarity matrices for real and estimated MV-patterns. The average coefficients between the lower triangular portions of the matrices are 0.19, 0.16 and 0.15 (respectively for ITC, FFA and PPA, all *p*<0.001). Visual inspections show that some patterns of the real dissimilarity matrices are preserved in the estimates (e.g. the patterns highlighted by the grey boxes for ITC and FFA).

**Fig. 5.**
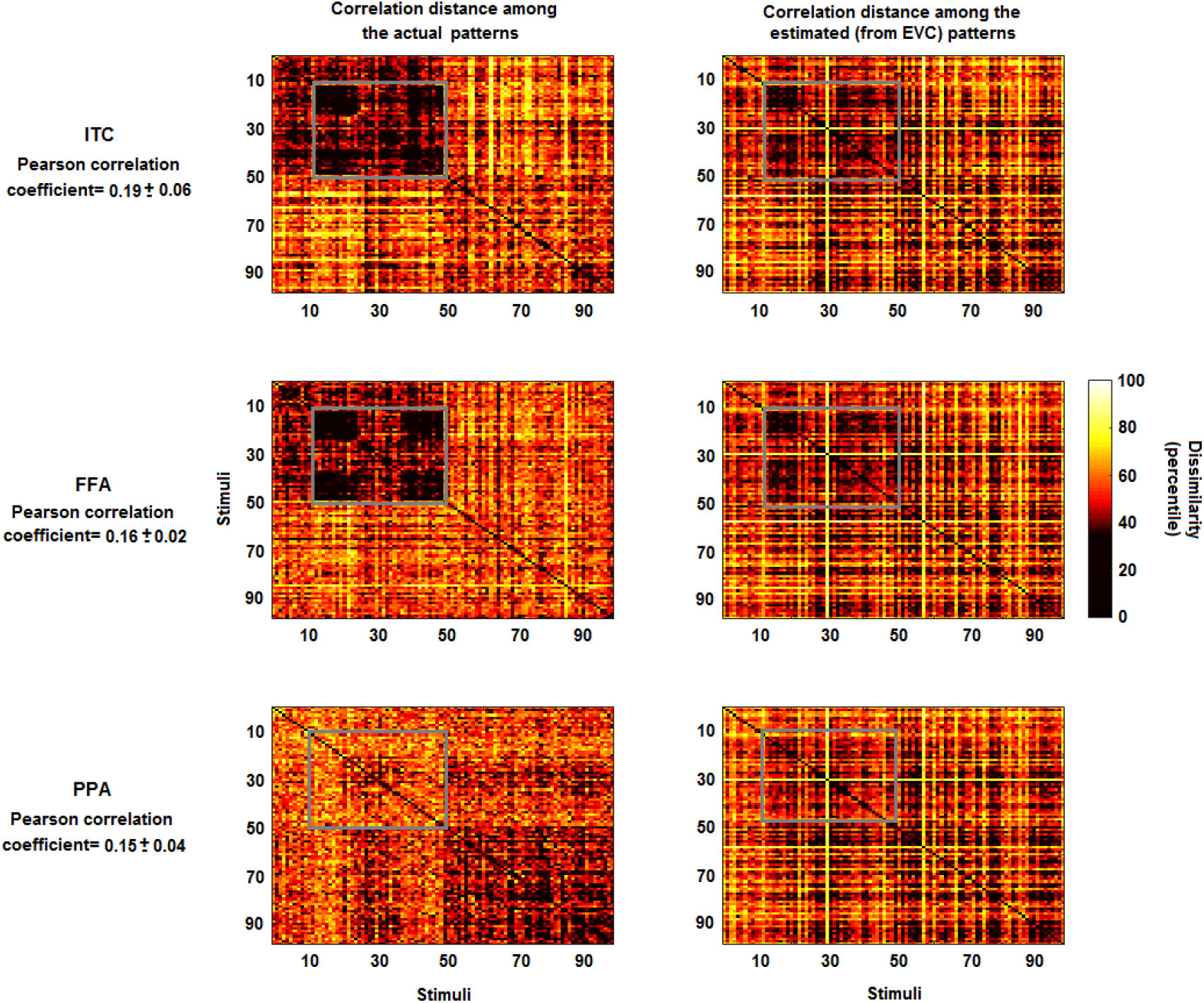
Representational dissimilarity matrices for actual and estimated patterns. Upper-middle- and lower left panels: percentile of the dissimilarity (computed by using the correlation distance) among the multivariate patterns of the actual ITC, FFA and PPA. Upper-middle- and lower right panels: percentile of the correlation distance among the estimated patterns from EVC. The estimates of the multivariate patterns were obtained by using the pattern transformation between EVC and ITC, FFA and PPA. While some information is lost, some characteristic patterns which are visible in the left panels (e.g. the patterns highlighted by the grey boxes for ITC and FFA) are also visible in the right panels. The average (across subjects) coefficients between the lower triangular portions of the representational dissimilarity matrices is 0.19, 0.16 and 0.15 (all p-values<0.001).

All the estimated regularization parameters λ lie within the range of [0.01, 10000], without reaching the boundaries of the interval. In particular, for the analysis on the 96 stimuli, the parameters for the transformations EVC-ITC were 1071.4±1 70.2, while for EVC-FFA and EVC-PPA were equal to 1573.6±334.7 and 2673.1±942.7. It is evident that there exists an anticorrelation between the *GOF* and λ values (Pearson correlation coefficient of −0.86, *p*<0.001). In particular, the higher the variance explained by the estimated transformation 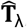, the lower the optimal value of the regularisation parameter λ.

Finally, we also investigated the GOF value of transformations obtained after the application of a k-fold cross-validation (with k=10), one other possible approach for defining the regularization parameter. The results were in accordance with those shown in Fig. 4.

### 3.2 Sparsity

Figure 6 shows the results obtained for the rate of decay of the density curve (*RDD*) and the *GOF* values (as described in the section 2.3.1), in order to characterise the sparsity of the MV-pattern transformations. We found a generally high level of sparsity for the transformations EVC->ITC, EVC->FFA and EVC->PPA. All the transformations reach an estimated sparsity >80% (Fig. 6). The transformation EVC->PPA (right panel, Fig. 6) shows the highest estimated levels of sparsity (>90%), followed by the transformation EVC->FFA (middle panel, Fig. 6) and EVC->ITC (left panel, Fig. 6) (both at 80-90%).

**Fig. 6.**
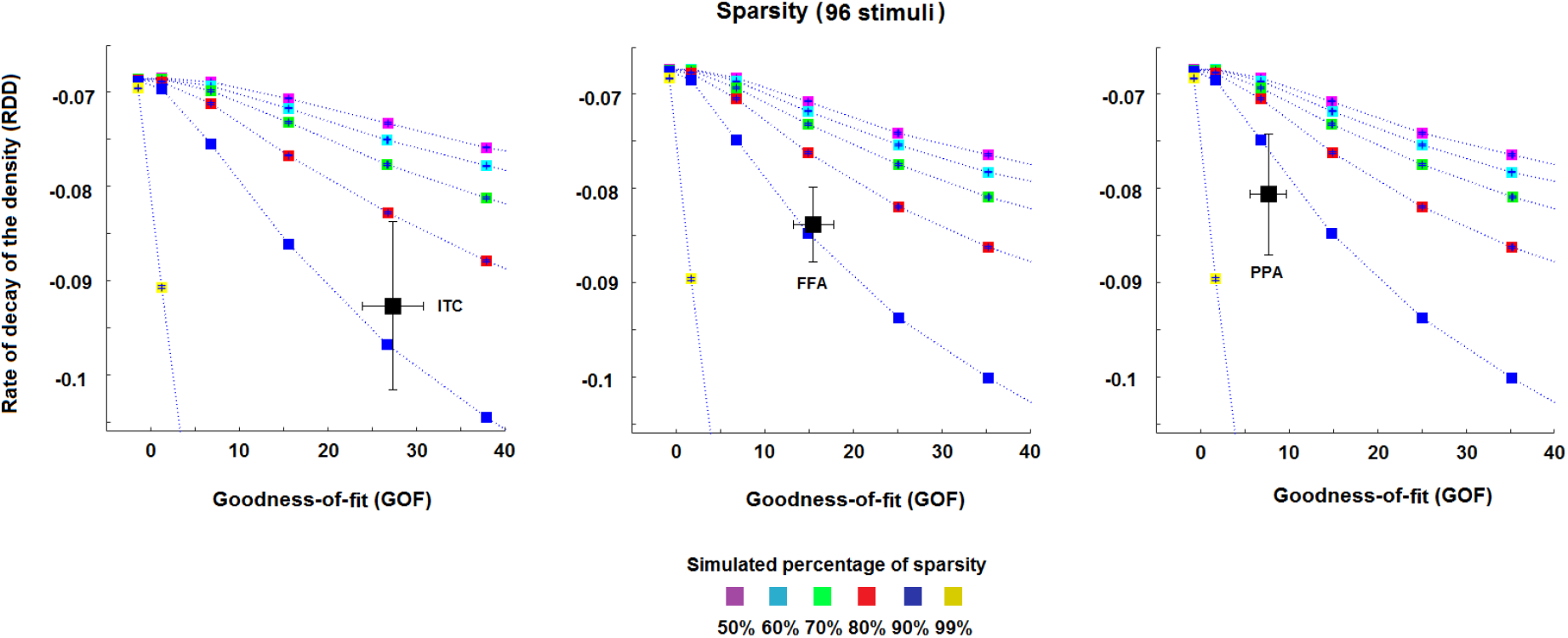
Estimated sparsity for the pattern transformation. The three panels show the estimates of the percentage of sparsity for the pattern transformations EVC->ITC, EVC->FFA and EVC->PPA, respectively. The dotted lines represent the mean *RDD* and *GOF* values across the simulationrealisations for each of the simulated percentages of sparsity, i.e. 50%, 60%, 70, 80%, 90% and 99%. Each area between two dotted lines thus represents a fixed range for the percentage of sparsity, e.g. the area between the two curves associated with the blues and red squares denotes a degree of sparsity between 80% and 90%. The black squares (and their error bars) in the panels denote the mean (and the standard error of the mean) estimate of *RDD* and *GOF* across the four subjects. All estimates show a high percentage of sparsity for all transformations (>80%). Slightly higher percentages are shown by the transformations EVC->FFA and EVC->PPA.

All estimates show large standard errors that, in some cases (see left panel, Fig. 6), may include curves associated with more than one percentage of sparsity. This suggests that an accurate estimate of the sparsity for the pattern transformations may benefit from a larger number of subjects and stimuli. Finally, the results in Fig. S5 clearly show that the average values for the session1-session2 mappings are in accordance with those obtained by considering the session2-session1 transformations, thus suggesting a within-subject stability of the metric.

### 3.3 Pattern deformation

Figure 7 shows the results for the pattern deformation metric, i.e. for the estimated rate of decay of the SVs (*RDSV*) and the *GOF* values (see section 2.4.1), for the transformations EVC->ITC, EVC->FFA and EVC->PPA. For the set of 96 stimuli, the transformations show different rates of decay of the SV-curve among them.

**Fig. 7.**
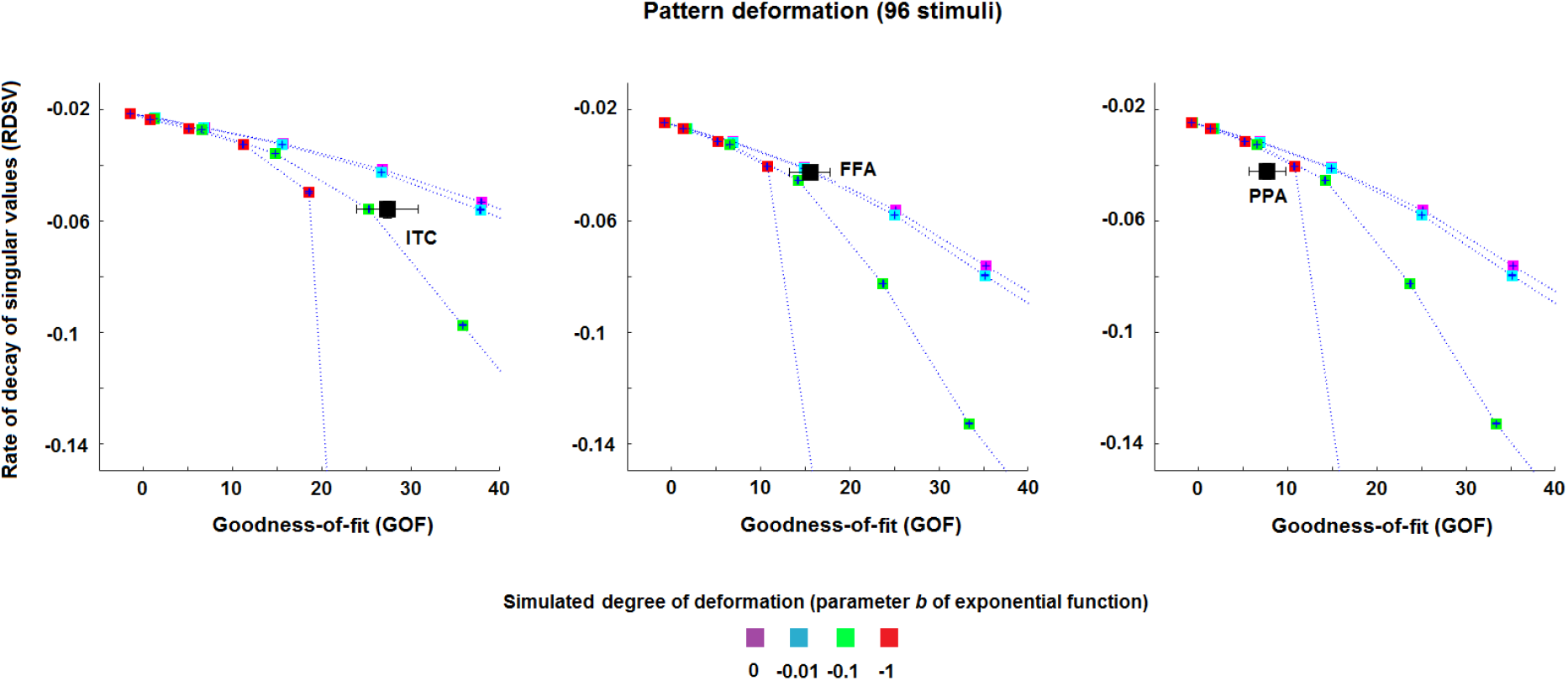
Pattern deformation for each pair of ROIs. The three panels show the estimates of the rate of decay of SV-curve (RDSV), denoting the pattern deformation, of EVC->ITC, EVC->FFA and EVC->PPA for the 96 stimuli. The dotted lines represent the mean *RDSV* and *GOF* values across the simulationrealisations for each of the simulated rates of decay. Each area between two dotted lines thus represents a fixed range for the rate of decay. The black squares (and their error bars) in the panels denote the mean (and the standard error of the mean) estimate of *RDSV* and *GOF* across the four subjects.

The results associated with the transformation EVC->FFA cover the curves associated with a deformation of −0.01 and 0 (these two curves practically coincide for low GOF values, middle panel of Fig. 7). This transformation is thus characterised by a more uniform deformation of the EVC MV-patterns than the other two transformations, which do not uniformly deform the patterns (the estimated parameter is approximately equal to or lower than −0.1 for EVC->ITC and EVC->PPA, respectively).

Although the error bars (standard error of the mean) show small values along the y-axis, the characterisation of the rate of decay of SV-curve would benefit from a larger number of subjects and stimuli, as well as from a higher signal-to-noise ratio. Finally, similarly to the sparsity metric, the results in Fig. S6 show that the average values for the session1-session2 mappings are in accordance with those obtained by considering the session2-session1 transformations.

## 4 Discussion

In this study, we developed computational strategies for estimating and analysing linear transformations between multivariate fMRI patterns in pairs of regions-of-interest (ROIs). These methods allow investigating features of the voxel-by-voxel mappings between ROIs. We first described a cross-validated ridge regression approach for robustly estimating the linear pattern transformation. Then, we described three novel metrics to characterise specific features of these transformations, i.e. the goodness-of-fit, the sparsity of the transformation and the pattern deformation. The first metric, goodness-of-fit, describes to what degree the transformations can be represented as a matrix multiplication, thus estimating the linear statistical dependency between two multi-voxel patterns. The second metric, sparsity, is closely related to the concept of topographic projections, i.e. possible one-to-one connections between voxels. The higher the percentage of sparsity, i.e. the higher the percentage of zero elements of the transformation, the higher is the degree to which the transformation represents a “one-to-one” mapping between voxels of the two ROIs. In order to estimate the percentage of sparsity, we relied on a Monte Carlo procedure to overcome the confounds induced by noise. The third metric, pattern deformation, is a measure of the degree to which the transformation amplifies or suppresses certain patterns. For instance, a constant value for the SVs of the transformation is associated with two multivariate patterns which can be seen as rotated versions of each other, while a larger decay is associated with a larger deformation (all the MATLAB functions and scripts for testing the metrics are freely available from https://github.com/alessiobastineuroscience/Analysing-linear-MV-pattern-transformation). We applied the ridge regression method, and the three different metrics, to an event-related fMRI data set consisting of data from four human subjects (Kriegeskorte et al. 2008a).

The results obtained using the goodness-of-fit measure showed the presence of a statistically significant linear dependency between EVC and the other three ROIs. Among the regions considered, ITC showed the highest linear dependency with EVC. Furthermore, in accordance with the existing literature (Kanwisher et al. 1997; Epstein & Kanwisher 1998; Mur et al. 2012), FFA showed the expected preference for faces and animate objects, while PPA for places and inanimate objects (Fig. 4 and S4). These findings indicate that, even if the true pattern transformations between brain areas might be non-linear, linear transformations can provide a good approximation. Importantly, while non-linear methods (such as neural networks) may increase the goodness-of-fit compared to linear methods (Anzellotti et al. 2016), linear methods allow the investigation of meaningful features of the transformation, such as sparsity and pattern deformation.

Our Monte Carlo approach for analysing sparsity revealed that our estimated linear transformations can be considered as sparse. Almost all the pattern transformations showed an estimated percentage of sparsity higher than 80%. Nevertheless, although the observed percentages suggest the presence of a one-to-few voxels mapping, they are lower than those expected for a precise one-to-one voxel mapping. Such a mapping between two ROIs, of e.g. 200 voxels each, would imply a percentage of sparsity higher than 99.5%. Importantly, the sparsity estimates reflect the percentage of components of the transformations that are equal to zero. These percentages, as opposed to the absolute number of zeros in the matrices, can be compared across connections (e.g. a percentage of sparsity of 80% in the transformation EVC-ITC could coincide with a percentage of 80% in the transformation EVC-FFA).

The results obtained by applying the pattern deformation metric showed that the transformations from EVC to ITC, FFA and PPA patterns are associated with different levels of deformations. The average deformation induced by the transformation from EVC to FFA was the one showing the most uniform amplification/compression, while the other two transformations are associated with higher degree of deformation. Thus, each of these transformations amplifies certain MV-patterns while dampening others. It is possible that this metric is related to the previous metric of sparsity. For example, sparser transformations may exhibit a lower degree of pattern deformation. This could be studies in the future using simulations and empirical data. However, both metrics still provide us with information on different types of features of the transformation. Here, we preferred to rely on two simulations that are as independent as possible.

Our results are based on data sets from only four subjects. This is not sufficient for a reliable statistical analysis, and these results should therefore be seen mostly as a proof-of-concept of our novel approach. However, the functional specificity of the patterns of goodness-of-fit and the high degree of sparsity, which suggests the presence of one-to-few voxels mappings, are promising hints that these methods will be useful for the characterisation of neural pattern transformations in future studies.

Several variations and extensions of our approach are possible. In order to estimate the pattern transformations, we relied on a ridge regression method (Hoerl & Kennard 1970). This method aims at minimising the *l*^2^ norm of the residuals as well as the *l*^2^ norm of the transformation itself. This is not the only approach that can be used as a regression analysis method. For example, one can also apply the least absolute shrinkage and selection operator (LASSO, Tibshirani 1996), which is a least-squares method with an *l*^1^ penalty term, or an elastic net approach, which contains a combination of both *l*^2^ and *l*^1^ penalties (Zou & Hastie 2005). However, these estimators may lead to sparse solutions even in the presence of non-sparse linear mappings (by using an elastic net approach all the estimated matrices show in our case a percentage higher than 90%, which is probably due to the presence of noise). This issue may be mitigated by combining our Monte Carlo method with those methodologies, reducing overestimates of sparsity by evaluating the transformations in settings that resemble the actual levels of noise in the data. Furthermore, classic algorithms (Boyd 2010) for solving these minimisation problems require the pattern transformation to be vectorised, and the input pattern to be transformed into a matrix composed of copies of the original pattern, thus requiring long computation times. Nevertheless, future work should compare pattern transformations estimated using different regression analysis approaches, and their effects on our novel transformation metrics. An advantage of our regression approach is that it produces explicit transformations, from which we can extract meaningful features. It remains to be seen if this is also the case for non-linear methods such as neural networks (Anzellotti et al. 2016) or multivariate kernel methods (O’Brien et al. 2016).

We found a high degree of sparsity for our estimated transformations, which is consistent with the presence of topographic mappings. In the future, one could define a metric for “pattern divergence” using information about spatial proximity of voxels, in order to test whether voxels that are close-by in the output region project to voxels that are also close-by in the input region.

Whereas in our work we explicitly focused on the well-known feedforward direction of information flow in the ventral visual stream, this clear relationship may not be evident for other ROIs. Moreover, there could also be a bidirectional interaction between two regions; in principle one could estimate the functional mapping from X to Y and from Y to X and analyse the features of both mappings in order to investigate the functionally relevant features of possible feed-back and feed-forward processes. Our method can also be applied to resting state data (where the transformation could be estimated using multivariate regression of the time courses in different regions as in Anzelotti et al. 2017, or Ito et al. 2017). It would then be possible to apply our metrics to these transformations. Our approach could also be generalized to the case of more than two ROIs. For instance, it would be possible to compute the transformation from pairs of regions while partialling out the contributions of other ROIs, thus leading to an estimate of the direct-transformation from the input to the output region.

Moreover, our method can be applied to other neuroimaging modalities, such as electro- and magneto-encephalography (EEG and MEG). This opens up the possibility of studying transformations across time, i.e. whether there are (non-)linear transformations that relate a pattern in an output region to patterns in an input region at different time points. While current approaches using RSA or decoding can test whether patterns or pattern similarities are stable over time (e.g. King & Dehaene 2014), our approach can potentially reveal whether there are stable or dynamic transformations among patterns of brain activity. In the linear case, this would be related to multivariate auto-regressive (MVAR) modelling (e.g. Stokes & Purdon 2017; Seth et al. 2015). So far, these methods have been used to detect the presence of significant connectivity among brain regions. Future work should investigate whether we can use the actual transformations to characterise the spatial structure of these connections in more detail. Our study demonstrates that linear methods can be a powerful tool in this endeavour, and may pave the way for more biophysically informed approaches using non-linear methods. Finally, our methods have translational potential. For example, they can be used to investigate whether the complexity of pattern transformations is affected by brain diseases such as dementias or schizophrenia, or how these transformations change across the life span.

## Supplementary material

**Fig. S1.**
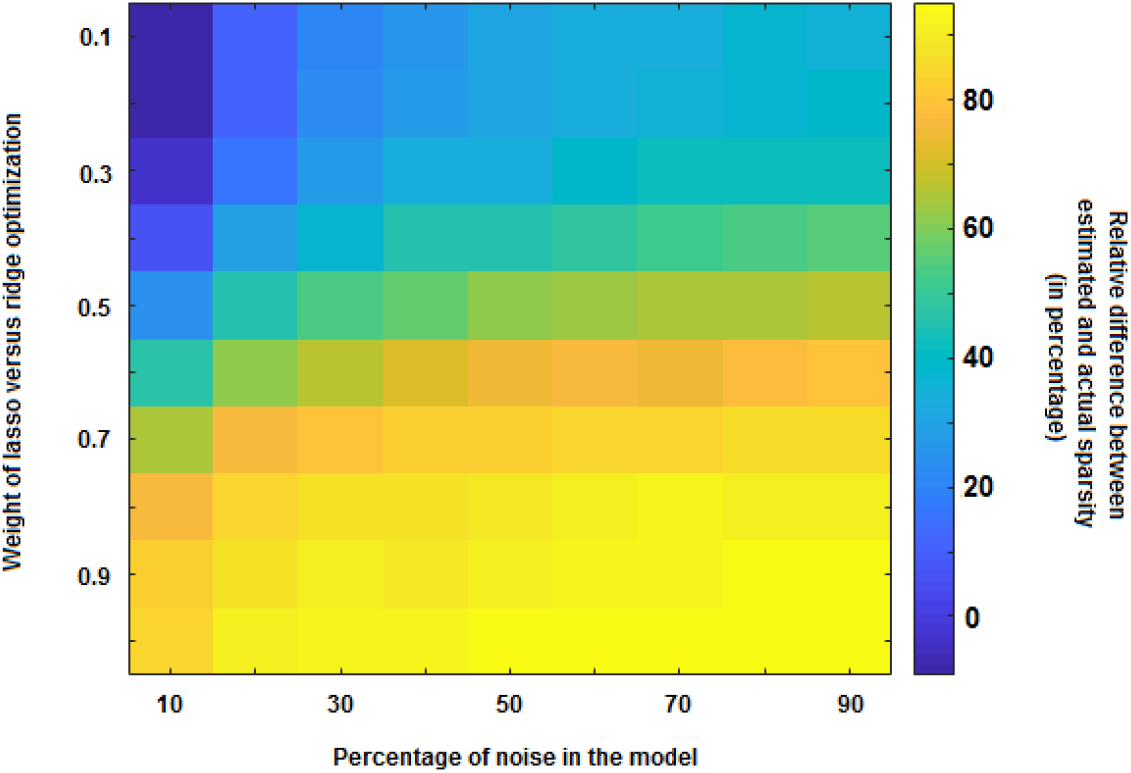
Percentage increment in estimating the simulated percentage of sparsity using different elastic net approaches. We simulated MV-interactions (with transformations between the two ROIs having a sparsity of 50%) and we estimated the mapping using different types of elastic net approaches, each of which consisted of a specific weighted combination of LASSO and ridge regression. The dimension of the simulated MV-pattern matrices was 224 (voxels) x 96 (stimuli) for the input ROI and 256 (voxels) x 96 (stimuli) for the output ROI, resembling the dimension of the actual early visual cortex (EVC) and fusiform face area (FFA) MV-pattern matrices. Regardless of the regularisation method, the estimated percentage of sparsity increases with increasing levels of noise. Furthermore, at low noise levels, the sparsity estimates improve slightly when the weight for ridge regression is higher than the weight for LASSO.

**Fig. S2.**
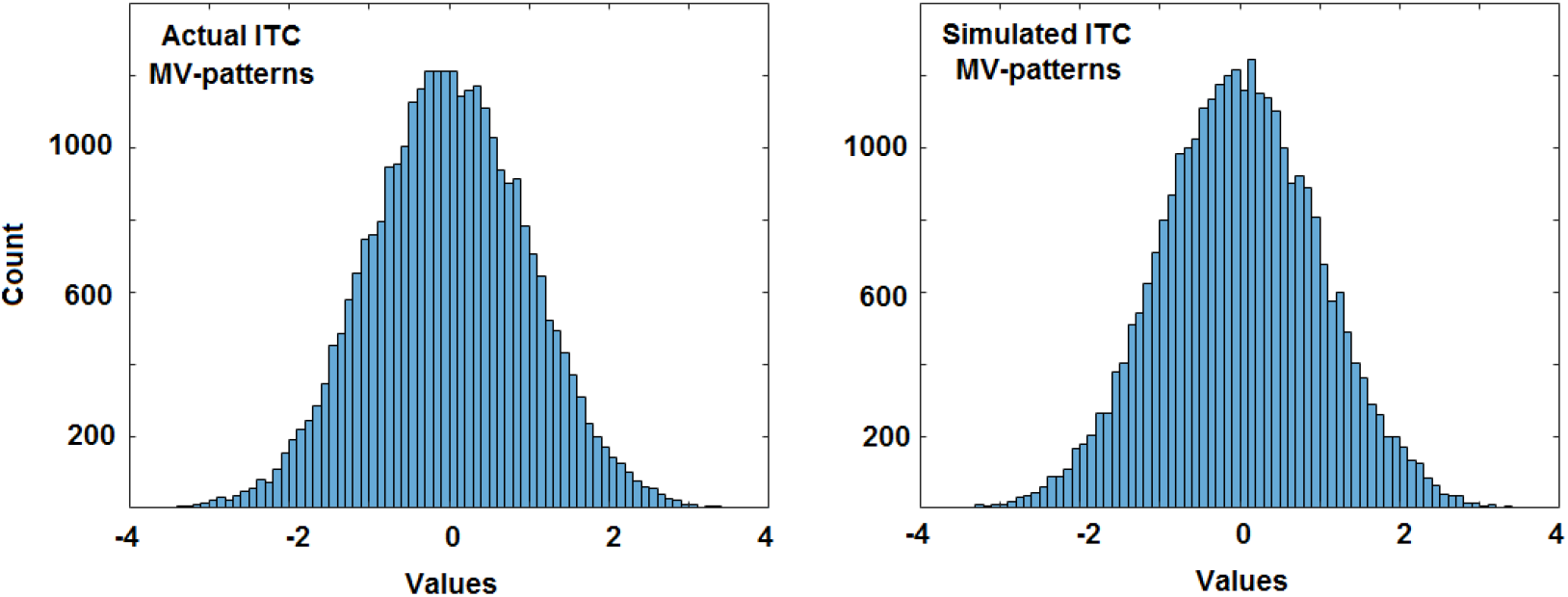
Histogram associated with the values of the actual (left panel) and simulated (right panel) MV-patterns of the inferior temporal cortex (ITC) for one subject. As for the Monte Carlo approach, the simulated ITC patterns have been generated as *Y* = (1 − *γ*)*TX*/||*TX*||_*F*_ + *γN*/||*N*||_*F*_ where denotes the EVC MV-pattern matrix for the subject, γ is a parameter representing the weight of the noise in the model (set as equal to 0.6 in this realisation), *N* is random noise and *T* is a simulated transformation (whose values follow a standard normal distribution).

**Fig. S3.**
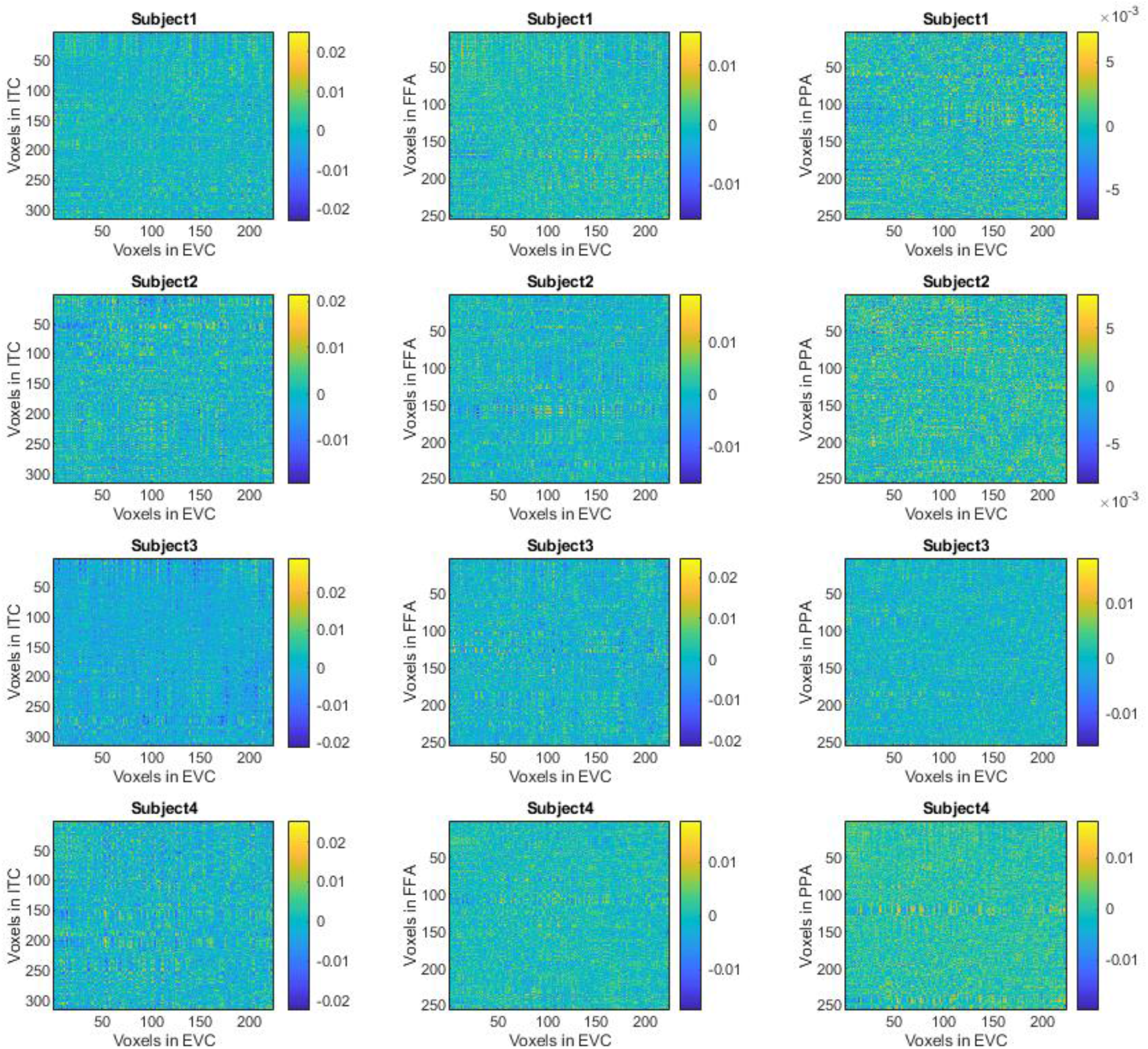
Estimated transformation for each subject and pair of ROIs. Each row shows the transformations between the MV-patterns of early visual cortex-EVC and the patterns of the other ROIs (inferior temporal cortex-ITC, fusiform face area-FFA and parahippocampal place area-PPA) for one subject. The columns refer to the specific pairs of ROIs (from left to right: EVC->ITC, EVC->FFA, EVC->PPA).

**Fig. S4.**
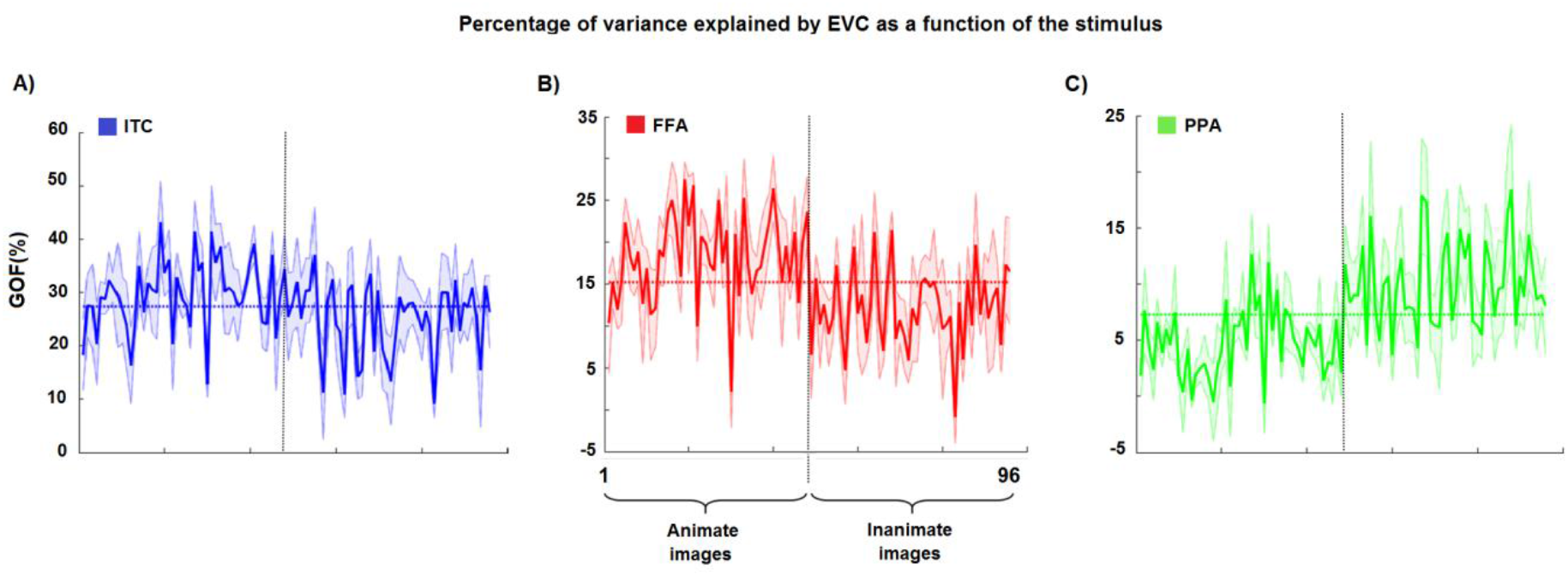
Goodness-of-fit (*GOF*) as a function of the stimuli. In order to compute the *GOF*, we here considered the squared Frobenius norm of the difference between the actual and estimated MV-pattern in the output ROI for each specific stimulus separately. **A)**, **B)** and **C)** The solid lines (and the shaded areas) denote the average percentages of *GOF* across the four subjects (and standard error), using the linear pattern transformations from EVC to the three output ROIs, as functions of the stimuli. The first 48 stimuli are animate images while the last 48 stimuli are inanimate images. A higher *GOF* (with respect to the mean) is evident for the animate stimuli for ITC and FFA, and for the inanimate stimuli for PPA.

**Fig. S5.**
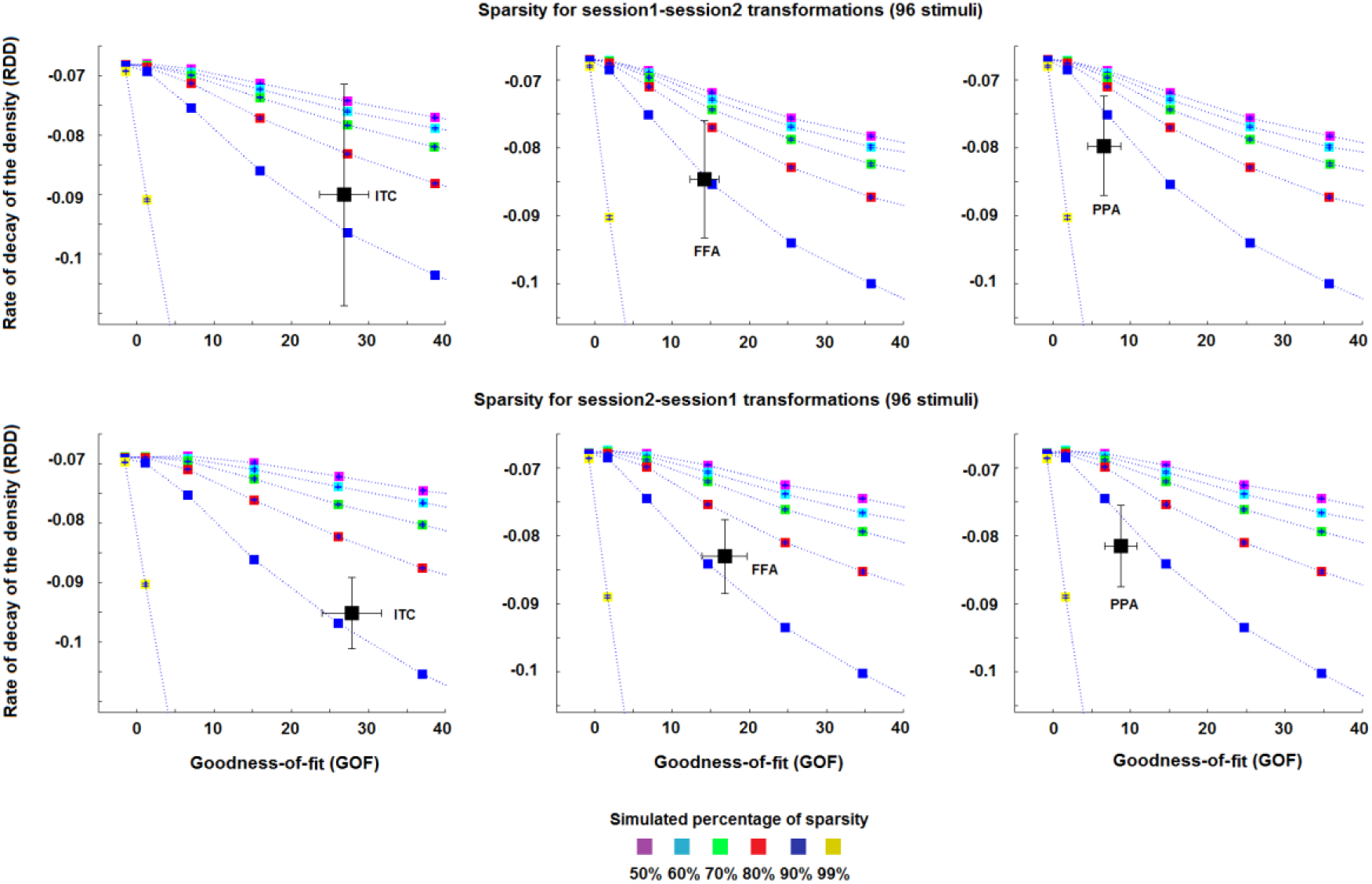
Estimated sparsity for the session1-session2 transformations and the session2-session1 transformations with 96 stimuli. The coloured squares (and their error bars) denote the mean (and the standard error of the mean) estimate of *RDD* and *GOF* across the 1000 simulation-realisations for each of the simulated percentages of sparsity, i.e. from 50% to 99%. The black squares (and their error bars) denote the mean (and the standard error of the mean) estimate of *RDD* and *GOF* across the four subjects. The results related to the session1-session2 transformations are in accordance with those obtained for the session2-session1 transformations.

**Fig. S6.**
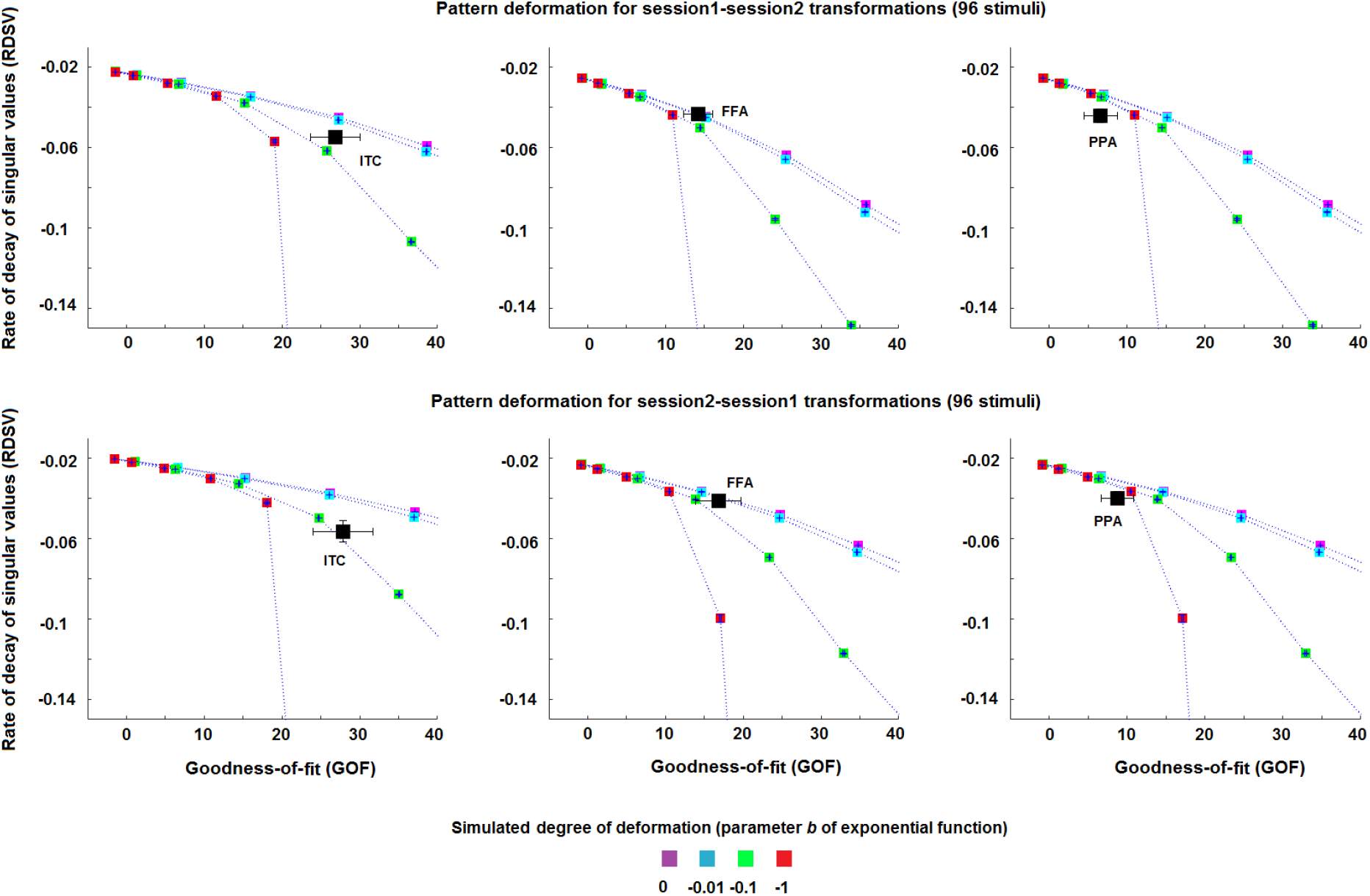
Estimated pattern deformation for the session1-session2 transformations and the session2-session1 transformations with 96 stimuli. The coloured squares (and their error bars) denote the mean (and the standard error of the mean) estimate of *RDSV* and *GOF* across the 1000 simulationrealisations for each of the simulated degree of deformation (parameter b of an exponential function). The black squares (and their error bars) denote the mean (and the standard error of the mean) estimate of *RDSV* and *GOF* across the four subjects. The results related to the session1-session2 transformations are in accordance with those obtained for the session2-session1 transformations.

